# Ultra-small cells and DPANN genome unveiled inside an extinct vent chimney

**DOI:** 10.1101/2021.03.08.430540

**Authors:** Hinako Takamiya, Mariko Kouduka, Hitoshi Furutani, Hiroki Mukai, Takushi Yamamoto, Shingo Kato, Yu Kodama, Naotaka Tomioka, Motoo Ito, Yohey Suzuki

## Abstract

Chemosynthetic organisms flourish around deep-sea hydrothermal vents where energy-rich fluids are emitted from metal sulfide chimneys. In contrast to actively venting chimneys, the nature of microbial life in extinct chimneys without fluid venting remains largely unknown. Here, the occurrence of ultra-small cells in silica-filled grain boundaries inside an extinct chimney is demonstrated by high-resolution bio-signature mapping. The ultra-small cells are associated with extracellularly precipitated Cu_2_O nanocrystals. Single-gene analysis shows that the chimney interior is dominated by a member of Pacearchaeota known as a major phylum of DPANN. Genome-resolved metagenomic analysis reveals that the chimney Pacearchaeota member is equipped with a nearly full set of genes for fermentation-based energy generation from nucleic acids, in contrast to previously characterized Pacearchaeota members lacking many genes for nucleic acid fermentation. We infer that the ultra-small cells associated with silica and extracellular Cu_2_O nanocrystals in the grain boundaries are Pacearchaeota, on the basis of the experimentally demonstrated capability of silica to concentrate nucleic acids from seawater and the presence of Cu-exporting genes in a reconstructed Pacearchaeota genome. Given the existence of ~3-billion-year-old submarine hydrothermally deposited silica, proliferation of microbial life using silica-bound nucleic acids might be relevant to the primitive vent biosphere.

## Introduction

The oldest record of pillow lava basalt, a geologic evidence of deep-sea hydrothermal activity, is found in the 3.8-billion-year-old Isua Supracrustal Belt in Greenland^1^. Deep-sea hydrothermal fluid venting by “black smokers” is associated with the extensive eruption of pillow lava on the deep seafloor^2^. Seawater is recharged, interacts with extrusive and intrusive basaltic rocks, and is then discharged as high-temperature hydrothermal fluid enriched with heavy metals and chemicals that can be used for microbial energy generation, such as HS^-^, Fe(II), CH_4_, and H_2_. As abiotic synthesis of nucleic acids and amino acids appears to be favorable from the fluid-borne chemicals, deep-sea hydrothermal vents are considered to be one of the potential habitats for primitive cellular life on Earth^3^.

As a result of rapid cooling of hydrothermal fluid, vent chimneys tend to form with an internal zonation of mineral assemblages^4^. Decreasing temperature and increasing pH cause the sequential deposition of minerals. In general, chalcopyrite (CuFeS2) precipitated from high-temperatures fluid (>300°C) tends to form the inner massive wall, which is surrounded by marcacite (FeS2), pyrite (FeS2), and sphalerite (ZnS) precipitated from lower temperatures in outer porous layers. Since the discovery of deep-sea hydrothermal vents^5^, microbial populations thriving in actively venting chimneys have been studied intensively^6^. However, in contrast to active chimneys with excess energy supplies from hydrothermal fluids, extinct chimneys without fluid venting appear to be hostile for chemolithotrophic microbes with scarce energy supplies available solely from metal sulfide minerals.

## Results and Discussion

### Extinct chimney inner wall with microbial signals

The Pika site is a recently discovered deep-sea hydrothermal field with black smokers (> 300°C) in the southern Mariana Trough, about 140 km east of Guam (**Figure 1a**)^7^. Chimney samples were collected by the remotely operated vehicle Hyper-Dolphin at a water depth and temperature of 2787 m and 1.7°C, respectively (**Figure 1b**). Zonation characteristic of metal sulfide chimneys was found in a thin section of one of the chimney samples (**Figure 1c**). There was an unaltered gold-colored part on the inner wall of chalcopyrite directly deposited from black smokers (**Figure 1d**). The chimney is referred to as IPdc in this manuscript.

**Figure 1.**
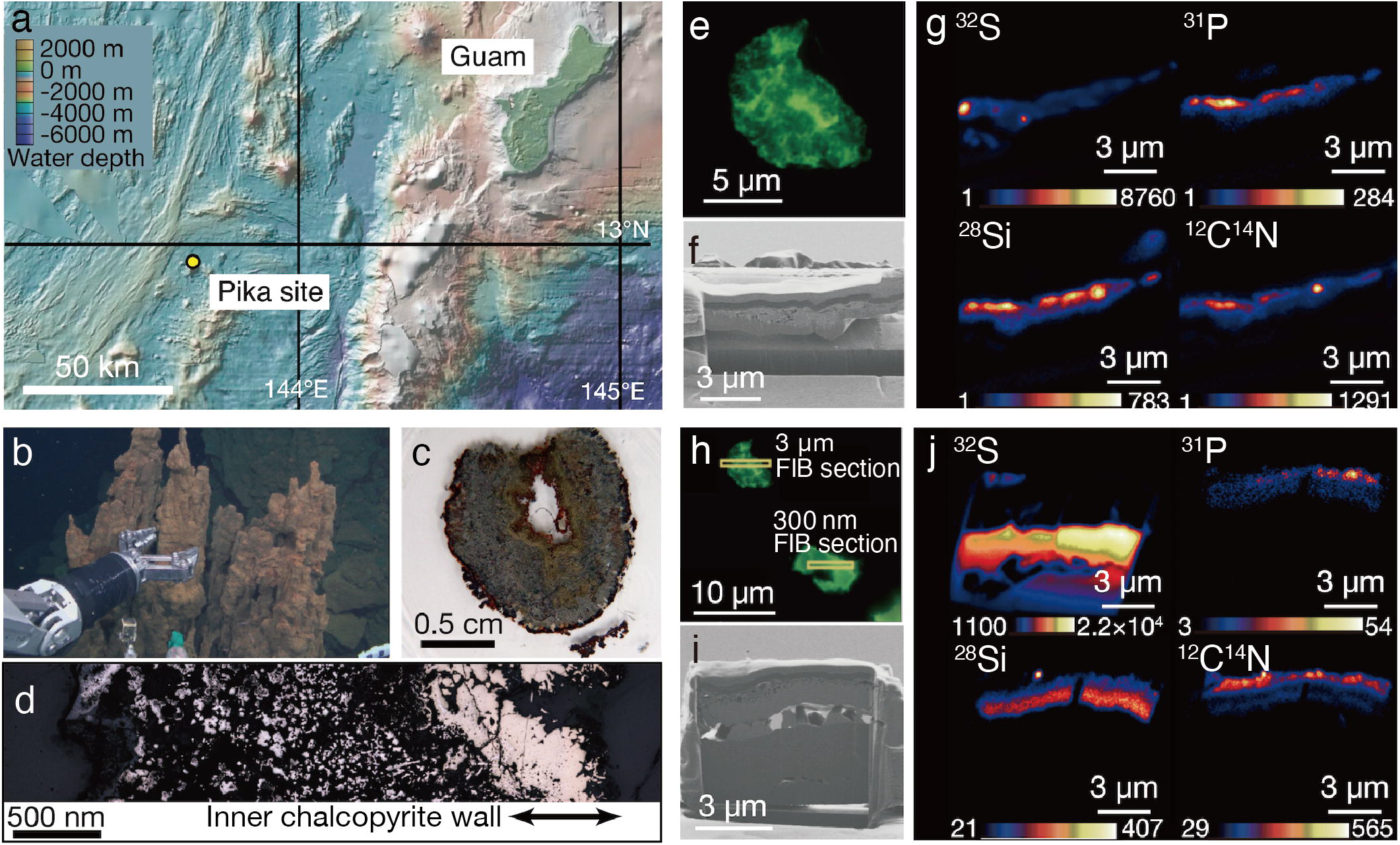
Sampling of an extinct chimney and microbial signal detections. **a,** Bathymetric map of the southern Marina Trough showing the Pika site and Guam. **b,** Photograph showing the extinct chimney sampled by the remotely operated vehicle *Hyper Dolphin*. **c,** Light microscopy image of a thin section from the extinct chimney. **d,** Reflection light microscope image of the thin section. The inner chalcopyrite wall is indicated with arrows. **e,** Fluorescence microscopy image of a grain boundary with greenish cell-like signals where a 3-µm thick focused ion beam (FIB) section was fabricated. **f,** Ga ion image of a 3-µm thick FIB section from the grain boundary shown in **Figure 1e. g**, Nanoscale secondary ion mass spectrometry (NanoSIMS) images of the FIB-fabricated grain boundary with intensity color contours. **h,** Fluorescence microscopy image of a grain boundary with greenish cell-like signals where a 300-nm thick FIB section was fabricated. **i,** Ga ion image of a 300-nm thick FIB section as pointed in **Figure 1i. j**, NanoSIMS images of the FIB-fabricated grain boundary with intensity color contours.

Analytical procedures have been developed to visualize and quantify microbial cells hosted in crack-infilling minerals in the oceanic crust by preparing thin sections of rocks embedded in hydrophilic resin called LR White^8,9^. Microbial cells embedded in the resin are stainable with a DNA dye called SYBR-Green I. Microscopic examinations of the inner wall of the chimney prepared as described above revealed the presence of patches with cell-like fluorescent signals where visible light was not transmitted (**Supplementary Figure 1**). Focused ion beam (FIB) sections with thicknesses of 3 μm (**Figure 1e,f,g**) and 300 nm (**Figure 1h, i, j**) were fabricated to characterize patches associated with the cell-like signals by nanoscale secondary ion mass spectrometry (NanoSIMS). In the 3-μm thick FIB section, signals from ^32^S, ^12^C^14^N, and ^31^P were detected in a silica-bearing layer with sub-micron voids (**Figure 1g, Supplementary Figure 2**). Similar results were obtained from the 300-nm thick FIB section (**Figure 1j**). A selected area electron diffraction (SAED) pattern showed that the silica-bearing layer also observed by transmission electron microscopy (TEM) was composed of amorphous material (**Supplementary Figure 3**).

### Microbial cells visualized in chalcopyrite grain boundaries

The greenish cell-like signals from the DNA dye and the overlapped NanoSIMS mapping of P and CN are consistent with results from previous study of microbial cells in mineral-filled cracks^9^. However, the appearance of individual cells was not clear, in contrast to the earlier work. To clearly observe individual cells, a 150-nm thick FIB section was fabricated from a chalcopyrite grain boundary associated with silica (**Supplementary Figure 4**) where greenish cell-like signals without light transmission were observed after SYBR-Green I staining (**Figure 2a**). It was revealed by NanoSIMS analysis that ^12^C^14^N, ^28^Si, and ^31^P were overlapped in a ribbon-shaped grain boundary (**Figure 2b, c**). TEM observations of the region with overlapping ^31^P, ^12^C^14^N, and ^28^Si showed small spheres with diameters of <200 nm (**Figure 2d, e**). Cuprite (Cu_2_O) was identified by a SAED pattern (**Figure 2f**) and an energy dispersive X-ray spectrum (EDS) (**Figure 2g**). High-resolution TEM observations revealed that ~5-nm-diameter particles of cuprite were spatially associated with the small spheres (**Figure 2h**).

**Figure 2.**
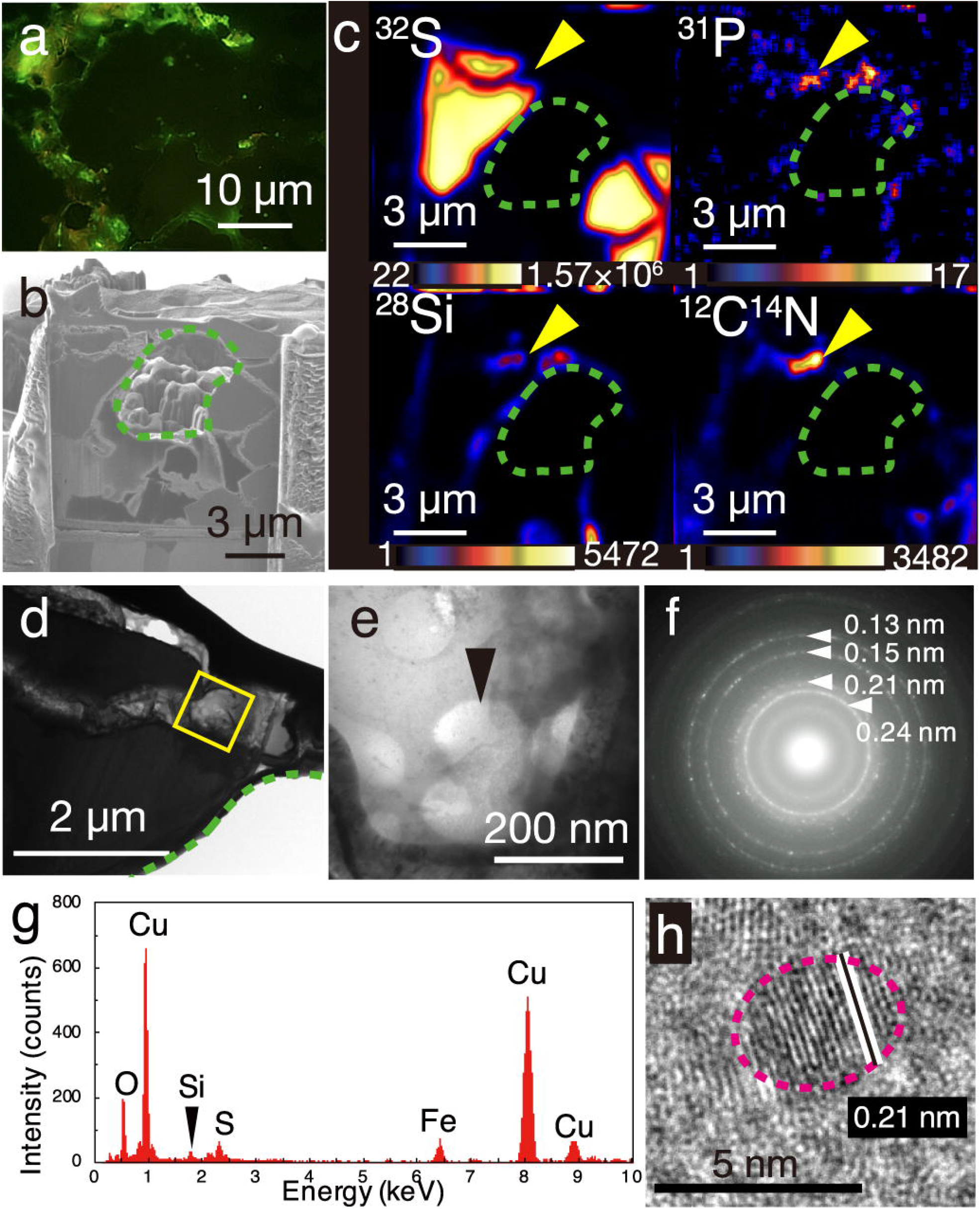
Visualization of microbial cells associated with Cu-bearing nanocrystals in chalcopyrite grain boundaries. **a,** Fluorescence microscopy image of a grain boundary with greenish cell-like signals where a 150-nm thick FIB section was fabricated. **b,** Ga ion image of a 150-nm thick FIB section of the grain boundary shown in **Figure 2a**. A green dotted line shows a hole corresponding to a chalcopyrite grain in the FIB section. **c,** NanoSIMS images of the 150-nm thick FIB section with intensity color contours. Yellow arrows indicate the region presented in Figure 2e. **d,** TEM image of a ribbon-shaped grain boundary of a 150-nm thick FIB section. A yellow square indicates the same region presented in Figure 2e. **e,** TEM image of small spheres. **f,** Selected area electron diffraction pattern from the small spheres in Figure 2e. Arrows indicate ring patterns with interplanar spacings. **g,** Energy dispersive X-ray spectrum from the small spheres. **h,** High-resolution TEM image of crystalline nanoparticles in the region pointed by a black arrow in Figure 2e. White lines indicate lattice fringes of a nanocrystal with the rim indicated by a pink dotted line.

To help interpret the TEM image of the small spheres associated with cuprite nanoparticles (**Figure 2e, Supplementary Figure 5a**), TEM observations were performed for cultured cells of *Geobacter sulfureducens* associated with extracellularly precipitated nanocrystals of uraninite (UO_2_) (**Supplementary Figure 5b, c**) and those of *Desulfovibrio desulfruicans* with UO_2_ nanoparticles in the periplasmic space (**Supplementary Figure 5d, e, f**). These observations clarified that the dark contrast is derived from the presence of uraninite nanoparticles, with transparent contrast from cellular materials and resin. The small spheres associated with cuprite nanoparticles in the chimney sample appear to be very similar to microbial cells associated with extracellular uraninite precipitation (**Supplementary Figure 5b**)^10,11^. The image contrast of the small spheres observed in the 150-nm thick FIB section was not transparent (**Figure 2e, Supplementary Figure 5a**). This is explained by the effect of small cell size. In sections with a thickness of 150 nm, transparent contrast is expected for cells with large cells, regardless of the cell shape (**Supplementary Figure 5g**). However, small coccoid cells are visualized with slightly dark contrast (**Supplementary Figure 5g**). The image contrast of the small spheres with extracellular Cu_2_O nanoparticles is therefore inferred to be derived from coccoid cells with a small size range. Small greenish spots visualized after DNA staining and the overlapped enrichment of P and CN revealed by NanoSIMS analysis support the conclusion that the small spheres are microbial cells.

### *In situ* biosignature analyses

To strengthen the evidence that the small spheres are indeed microbial cells, we performed *in situ* biosignature analyses. The spatial distributions of biomolecules such as proteins and lipids in chalcopyrite grain boundaries were characterized by an imaging mass spectrometry using a thin section after SYBR-Green I staining and observations by fluorescence microscopy. Spot analysis of grain boundaries revealed the presence of a macromolecule with *m/z* = 805.2 (**Supplementary Figure 6**). The mass spectrum of resin was found to lack this macromolecule. Mapping of the macromolecule in the chimney inner wall confirmed its ubiquitous distribution in grain boundaries (**Figure 3a, b**).

**Figure 3.**
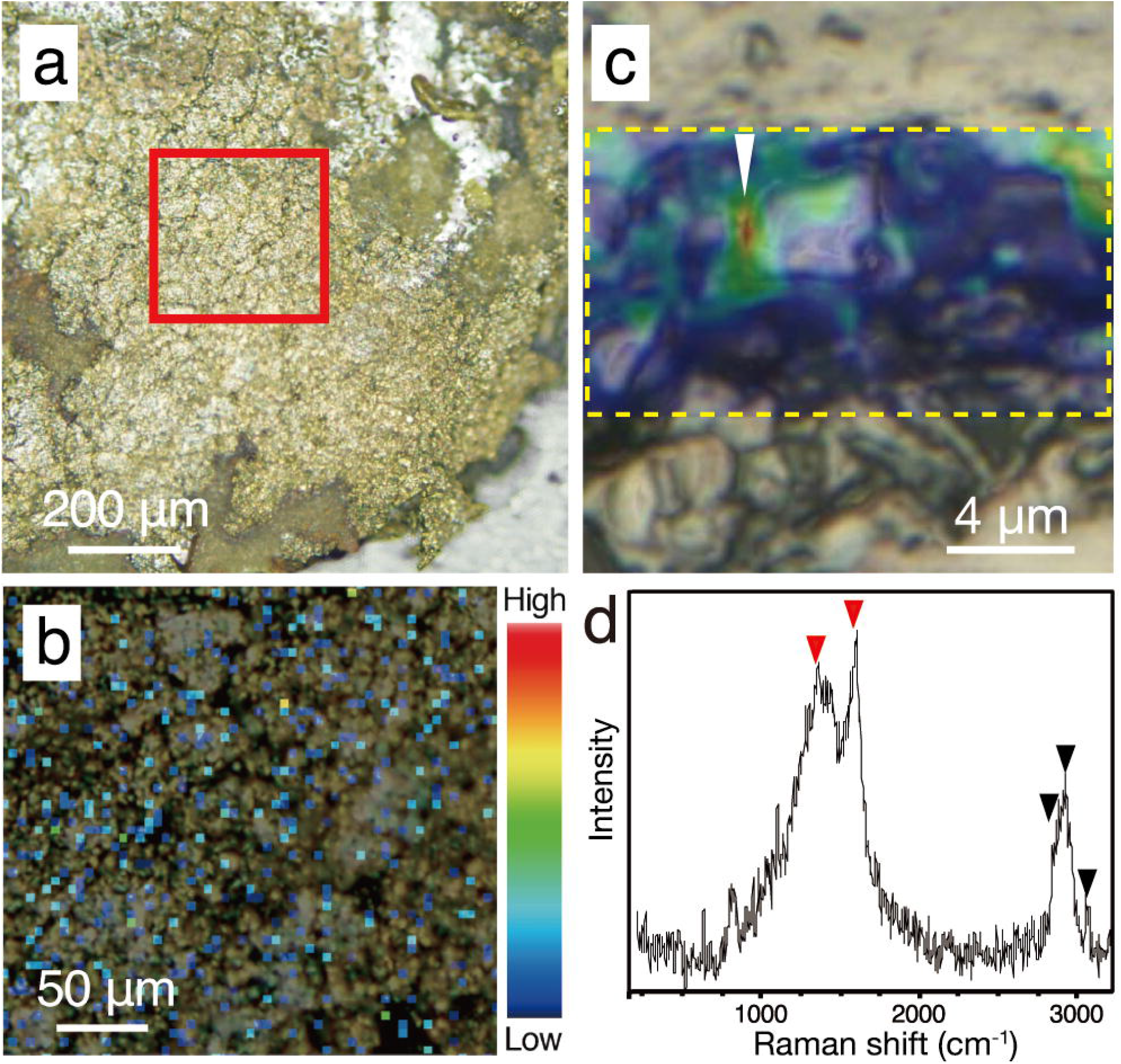
Biological signatures determined by imaging mass spectrometry and Raman spectroscopy from chalcopyrite grain boundaries. **a,** Reflection light microscope image of inner chalcopyrite wall. The red square highlights the region observed at a higher magnification in **Figure 3b. b**, Mapping image of a macromolecule at with *m/z* =805.2 by imaging mass spectrometry analysis with an intensity color contour. **c,** Optical microscope image overlapped with a Raman spectroscopy mapping image of the peak intensity at 2800–3000 cm^-1^, as highlighted by a yellow dotted rectangle. **d,** Raman spectrum obtained from the region indicated by a white arrow in **Figure 3c**. Red arrows indicate peaks not found in the resin or minerals spatially associated with the small spheres, while back arrows indicate peaks also found in the resin (**Supplementary Figure 7**).

To obtain another line of convincing evidence for the biogenicity of the small spheres^12^, Raman spectroscopy was used to characterize the same region where NanoSIMS analysis was performed on the ribbon-shaped grain boundary (**Figure 2c**). High-resolution mapping was performed for the peak intensity at 2800–3000 cm^-1^, which is attributed to CH_2–3_ typically found in microbial lipids (**Figure 3c**)^13^. The peak intensity was strong along grain boundaries. Raman spectra obtained from the grain boundaries with the high CH_2-3_ signal were different from those of the resin and minerals spatially associated with the small spheres (**Figure 3d, Supplementary Figure 7**). The main spectral difference is explained by the presence of CH3 and amide groups (C–N and N–H; indicated by red arrows), which is consistent with the distribution of ^12^C^14^N determined by NanoSIMS analysis (**Figure 2c**). We exclude the possibility that the small spheres are abiotic carbonaceous particles such as recently discovered particulate graphite (C) in hot and cold vent fluids^14^. In addition to the circularity and narrow size distribution of the spheres, the lack of sharp edges typically found in abiotic carbonaceous particles further supports the biogenicity of the small spheres^15^. Taken together, we conclude that the small spheres are ultra-small cells.

### Microbial abundance and composition in the extinct chimney

To understand microbial life in the chalcopyrite inner wall, the interior and exterior of the extinct chimney was carefully separated for cell counting and 16S rRNA gene amplicon analysis in combination with bulk mineralogical analysis. Cell densities in the chimney interior and exterior were determined by microscopic observations to be 1.9×10^8^ and 1.4×10^8^ cells/cm^3^, respectively, similar to those in extinct chimneys from Central Indian Ridge and western Pacific^16^. Single-gene analysis based on 16S rRNA gene sequences revealed that members of Pacearchaeota formerly referred to as Deep Sea Hydrothermal Vent Euryarchaeotal Group 6 (DHVE-6) were predominant in the chimney interior, remarkably different from the chimney exterior, which was dominated by members of the bacterial phylum Nitrospirae (**Supplementary Figure 8 and Supplementary Table 1**). Powder X-ray diffraction (XRD) analysis revealed both subsamples were composed of chalcopyrite, marcasite and pyrite (**Supplementary Figure 9**). It is likely that the remarkable difference in microbial composition can be attributed to textural features of the chimney interior (massive) and exterior (porous) (**Figure 1d**). This notion is supported by very minor occurrences of Pacearchaeota in previous studies of porous parts of extinct chimneys mainly containing chalcopyrite^16,17^. The absence of Epsilonproteobacteria, indicator organisms of active fluid venting in deep-sea hydrothermal fields^17^, is consistent with the extinct nature of the chimney sample.

### Genomic features of the chimney Pacearchaeota

As 16S rRNA gene amplicon analysis supported the idea that the Pacearchaeota members are predominant in the inner chalcopyrite wall, a near-complete genome of Pacearchaeota representative of 16S rRNA gene sequences was reconstructed by metagenomic analysis. In addition to analysis of a 16 rRNA gene sequence in the reconstructed genome (**Figure 4a**), ribosomal protein S3 region (rpS3) gene sequence analysis further confirmed phylogenetically affiliation with the Pacearchaeota phylum (**Figure 4b**). The genome size was very small (0.81 Mbp), with a genomic completeness of 90.4%, based on 52 single copy genes (**Supplementary Tables 2, 3**) ^18^. The small genome is consistent with genome sizes of Pacearchaeota members from previous studies (**Figure 4c**)^18–20^.

**Figure 4.**
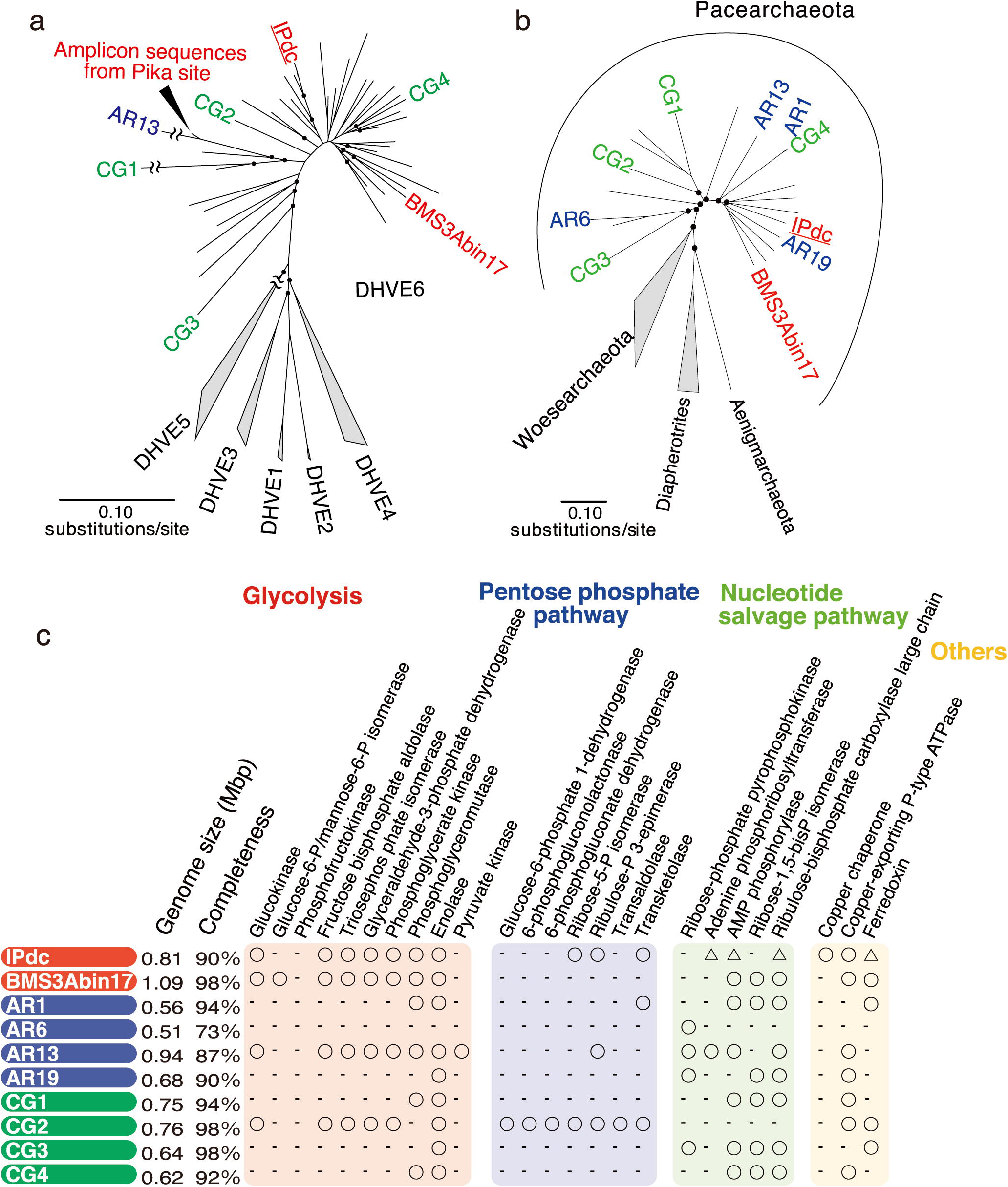
Comparative analysis of Paceacrhaeota-affiliated genomes from present and previous studies. **a–b,** Neighbor-joining phylogenetic trees based on nucleotide sequences of 16S rRNA (left) and amino-acid sequences of ribosomal protein S3 (right). Branches with bootstrap values >50% are indicated with filled circles. Colors indicate sampling sources: deep-sea metal sulfide deposits (red)^20^; shallow groundwater, Rifle Site, Colorado (blue)^18^; and deep groundwater, Crystal Geyser Site, Utah (green)^19^. **c,** Overview of presence or lack of genes for metabolic pathways such as glycolysis, the pentose phosphate pathway, and nucleotide salvage pathway. The lack of genes does not indicate their absolute absence in the original genome, because of the incompleteness of genome reconstruction. KEGG Orthology (KO) numbers for these genes are listed in Supplementary Table 5. A circle indicates the presence of the gene in the near-complete IPdc Pacearchaeota genome, whereas a triangle indicates its presence in contig sequences assembled from IPdc sequence fragments.

With respect to carbon metabolism, nearly full sets of genes involved in glycolysis and the non-oxidative pentose phosphate pathway (PPP) were encoded in the IPdc Pacearchaeota genome, indicating adenosine triphosphate (ATP) generation via fermentation of sugar-based biomolecules (**Figure 4c, Supplementary Figure 10, Supplementary Table 4**). Pacearchaeota-related genes for the nucleotide salvage pathway, including those encoding adenosine monophosphate (AMP) phosphorylase *(deoA)* and ribulose 1,5-bisphosphate carboxylase/oxygenase *(RuBisCO),* were found in IPdc contigs without being binned into genomes (**Figure 4c, Supplementary Table 5**). Phylogenetic analysis revealed that Pacearchaeota-related RuBisCO from IPdc contigs was classified as Type III-b and closely related RuBisCO genes in the Pacearchaeota genomes from the previous studies (**Supplementary Figure 11**)^18–20^. Amino acid sequence motifs that control catalytic activity and substrate binding were identical to those found in *Methanocaldococcus jannaschii*, the purified RuBisCO of which has carboxylase and oxygenase activities^21^.

Archaeal type III RuBisCOs are not associated with the typical role of RuBisCO in CO_2_ fixation in the Calvin-Benson-Bassham cycle; instead, they have recently been demonstrated to be involved in AMP metabolism^22,23^. In this pathway, AMP is converted to ribulose-1,5-bisphosphate (RuBP), and then the generation of two molecules of glycerate 3-phosphate from RuBP and CO_2_ is catalysed by the type III archaeal RuBisCOs (**Supplementary Figure 11**). These reactions are important to feed glycerate 3-phosphate into glycolysis for extra ATP generation.

### Cross-linking cellular and genomic features

We could not directly identify the ultra-small cells by performing fluorescence *in-situ* hybridization in thin sections (**Supplementary Figure 12**)^24^. Thus, we attempted to correlate the IPdc Pacearchaeota genome with the ultra-small cells in the grain boundary of the inner chalcopyrite wall. Given DPANN archaea with small cells have small genomes^25–27^, the <200-nm sized cells observed by TEM (**Figure 2e**) are consistent with the small genome size of the IPdc Pacearchaeota genome (**Figure 4c**). Cuprite nanoparticles, which were extracellularly associated with the ultra-small cells (**Figure 2f, g, h**), are thermodynamically stable in anoxic and neutral-to-alkaline conditions^28^. In the presence of O_2_, the oxidation of cuprite nanoparticles is fast^29^. These cuprite characteristics support the inference that the grain boundary has been anoxic since the termination of fluid venting. This inference is also supported by the presence of a Pacearchaeota-affiliated ferredoxin gene, which is an indicator of anaerobic metabolism^30^. The hyperthermopholic acraeon *Archaeoglobus fulgidus* is known to use Cu-exporting ATPase (CopA) and soluble Cu chaperone (CopZ) for translocating intracellular Cu(I) through cytoplasmic membrane (**Supplementary Figure 10**)^31^. Both genes encoded in the IPdc Pacearchaeota genome appear to play key roles in the extracellular formation of Cu(I)-bearing cuprite nanoparticles. As cuprite nanoparticles have anti-bacterial effects^32^, the close association with cuprite nanoparticles could help the Pacearchaeota member to dominate against bacterial competitors in the chimney interior.

Nanosolid analyses revealed that the grain boundaries of the inner chalcopyrite wall were associated with amorphous silica. Amorphous silica is known to adsorb nucleic acids and is widely used in molecular biology methods to purify DNA and RNA from other cellular components such as proteins and lipids^33^. We hypothesize that the ability of amorphous silica to selectively concentrate nucleic acids from seawater might explain the dominance of the Pacearchaeota member that produces ATP from nucleic acid fermentation in the grain boundaries (**Supplementary Figure 10**). To demonstrate binding of nucleic acids to amorphous silica in seawater, DNA extracted from *Escherichia coli* was loaded into a silica-based spin column. In slightly alkaline conditions (pH ~8), DNA was strongly bound to amorphous silica in seawater, whereas DNA binding was very limited in a low-salinity solution such as TE buffer (**Supplementary Figure 13**). Thus, the possibility that silica-bound nucleic acids could serve as carbon and energy sources in the chimney interior was experimentally confirmed.

### Implications

In the chimney inner wall, most of organic matter such as nucleic acids is initially destructed by black smoker (>300°C)^34^. Regardless of venting low- and high-temperature fluids, amorphous silica is ubiquitously formed inside the chimney by cooling of hydrothermal fluids^35^. By utilizing nucleic acids bound on amorphous silica from seawater, microorganisms could obtain sufficient energy and carbon sources to thrive inside extinct chimneys.

DPANN archaea are deep-branching and potentially diverged from non-DPANN archaea at an early evolutionary stage^36^. DPANN archaea are known to have small genome sizes and lack many biosynthetic pathways^18,37^. In particular, a member of Pacearchaeota has the smallest genome among DPANN archaea^37^. The Pacearchaeota members from deep-sea metal sulfide deposits have relatively many genes for glycolysis, the PPP and the nucleotide salvage pathway, whereas many Pacearchaeota members from groundwater samples lack many genes involved in nucleic acid fermentation in their reconstructed genomes^18,37^. We suggest that these genes are necessary for survival in the deep-sea rocky biosphere using nucleic acids bound to amorphous silica and/or released from other microorganisms.

The interior of the extinct chimney, found to be dominated by Pacearchaeota, appears to be environmentally similar to the early anoxic ocean on Earth. Salinity is an important factor controlling binding of nucleic acids to silica. The salinity range of the primitive ocean is estimated to be comparable to, or higher than, that of the modern ocean^38^. We therefore suggest that auxotrophy based on nucleic acid fermentation encoded in Pacearchaeota and other DPANN genomes may have played important roles in underpinning microbial ecosystems at hydrothermal vents on the early Earth. This possibility is supported by silica-bearing filaments within massive chalcopyrite textures in a 3.25-billion-year-old submarine hydrothermal deposit^39^.

## Methods

### Sample collection and subsampling

A metal sulfide chimney was collected from an active hydrothermal vent field at the Pika site (12°55.15’N, 143°36.96’E) in the Southern Mariana Trough during the Japan Agency for Marine-Earth Science and Technology (JAMSTEC) Scientific Cruises NT12-24 of the R/V *Natsushima* in September of 2012. The Southern Mariana Trough is a back-arc basin where the Philippine Sea Plate is subducted (**Figure 1a**). The chimney structure visually confirmed for the lack of hydrothermal fluid venting was collected by the manipulator arm of remotely operated vehicle (ROV) *Hyper Dolphin* (**Figure 1b**). The metal sulfide chimney was placed into an enclosed container in order to minimize contamination from surrounding seawater during transportation to the surface.

The collected chimney was immediately subsampled onboard in a cold room at 4°C. Using sterile chisels and spatulas, the exterior portion of the sample was subsampled. For subsampling of the interior portion, the chimney surface was flamed by a gas torch to prevent contamination from the exterior potion. Some intact portions of the chimney structure were preserved for light and electron microscopic observations, μ-Raman spectroscopy, imaging mass spectrometry and nanoscale secondary ion mass spectrometry (NanoSIMS) analysis for mineral and microbial distributions, while the rest of the chimney structure was ground into powder by using sterile pestles and mortars. Both the intact and the ground subsamples were fixed with 3.7% formamide in seawater onboard. The rest of the subsamples were frozen at −80°C for DNA extraction and mineralogical characterizations. All solutions were filtered with a 0.22-μm-pore-size filter (Millipore).

### Mineralogical and microbiological characterizations of thin sections from the metal sulfide chimney

To clarify mineral composition and microbial distribution within the chimney interior, thin sections were prepared using LR White resin. The intact chimney subsamples were dehydrated twice in 100% ethanol for 5 min, and then infiltrated four times with LR White resin (London Resin Co. Ltd.) for 30 min and solidified in an oven at 50°C for 48 h. Solidified blocks were trimmed into thin sections and polished with corundum powder and diamond paste. For the staining of microbial cells embedded in LR White resin, TE buffer with SYBR Green I (TaKaRa-Bio, Inc.) was mounted on thin sections and covered with cover glasses. After dark incubation for 5 min, thin sections rinsed with deionized water and mounted with the antifade reagent VECTASHIELD (Vector Laboratories) were observed using a fluorescence microscope (Olympus BX51) and a charge-coupled device (CCD) camera (Olympus DP71). Two ranges of fluorescence between 540–570 nm and 570–600 nm were used to discriminate microbial cells from mineral-specific fluorescence signals.

Mineral assemblages and textures were observed using the same microscope with the transmission light mode. Reflection light microscopy observations were conducted by an optical microscope (Nikon ECLIPSE E600POL E6TP-M61) and a CCD camera (Zeiss AxioCam MRc 5). Carbon-coated thin sections were characterized using a scanning electron microscope (Hitachi S4500) at an accelerating voltage of 15 kV. Back-scattered electron imaging coupled to energy-dispersive X-ray spectroscopy (EDS) was used to analyze the chemical compositions of mineral phases according to image contrasts.

To analyze microbial cells found in thin sections by NanoSIMS at the Kochi Institute for Core Sample Research (KOCHI), JAMSTEC (CAMECA NanoSIMS 50L), 3-µm- 300-nm- and 150-nm-thick sections were fabricated using focused ion beam (FIB) sample-preparation and micro-sampling techniques using a Hitachi FB-2100 instrument at the University of Tokyo or a Hitachi SMI-4050 at KOCHI. The thin-section samples were locally coated with the deposition of W (100–500-nm thick) for protection and trimmed using a Ga-ion beam at an accelerating voltage of 30 kV. A focused primary Cs^+^ ion beam of approximately 1.0 pA (100-nm beam diameter) was rastered on the samples. Secondary ions of ^12^C^−^, ^16^O^−^, ^12^C_2_^−^, ^12^C^14^N^−^, ^28^Si^−^, ^31^P^−^ and ^32^S^−^ were acquired simultaneously with multidetection using seven electron multipliers at a mass-resolving power of approximately 4500. Each run was initiated after stabilization of the secondary ion-beam intensity following presputtering of < approximately 2 min with a relatively strong primary ion-beam current (approximately 20 pA). Each imaging run was repeatedly scanned (15 to 20 times) over the same area, with individual images comprising 256 × 256 pixels. The dwell times were 5,000 μs/pixel for the measurements, and total acquisition time was approximately 2 h. The images were processed using the NASA JSC imaging software for NanoSIMS developed by the Interactive Data Language program^40^.

Transmission electron microscopy (TEM) was used to examine microbial distributions and the structure and composition of minerals at the nanometre scale. JEOL JEM-2010 with energy dispersive X-ray spectrometry (EDS) was operated at 200 kV at the University of Tokyo. A JEOL JEM-ARM200F transmission electron microscope was used at an accelerating voltage of 200 kV at KOCHI, JAMSTEC.

### Imaging mass spectrometry (MS)

An atmospheric pressure matrix-assisted laser desorption/ionization system equipped with a quadrupole ion trap-time of flight analyzer (MALDI-TOF) was used to obtain MS data from spots or regions observed by microscopy (iMScope *TRIO,* Shimadzu). The thin section subjected to nanosolid characterizations was further thinned by a mechanical polisher (Leica EM TXP Target Preparation Device). The thin section was coated with 9-aminoacridine (Merck) by iMLayer (Shimadzu) with a thickness of 1.0 μm and then irradiated by a 355 nm Nd:YAG laser with a laser diameter of ~5 μm. The positive ion mode was used for imaging of the thin section. Scanning was performed with a pitch of 5 μm. Operation conditions were as follows: frequency, 1000 Hz; 50 shots per spot. To visualize the ion images, Imaging MS Solution (Shimadzu) was used. For a reference, LR White resin sectioned into a thickness of 20 μm by the ultramicrotome was mounted on a carbon tape.

### Raman spectroscopy

High-resolution confocal Raman system (Horiba LabRam HR) equipped with a laser (532 nm) was used to characterize chalcopyrite grain boundaries on a thin section. The incident laser was operated at 1.2–4 mW. Analytical uncertainty in the Raman shift was ~2 cm^-1^, and the spatial resolution was ~1 μm. Raman spectra were compared with standard spectra obtained from RRUFF (http://rruff.info), and peak assignments were based on references^13,41–43^.

### Uranium reduction experiments and TEM observations of incubated cells

*Desulfovibrio desulfuricans* [ATCC#642] and *Geobacter sulfurreducens* [ATCC#51573] were subjected to uranium reduction experiments previously performed for *Desulfosporosinsus* spp.^44^ In an anaerobic glove box with a gas mixture of N_2_-CO_2_-H_2_ (90:5:5), cells enriched in media recommended by ATCC were incubated in 0.25% bicarbonate solution at pH 7 containing 1 mM of uranyl acetate. 10 mM sodium lactate and 10 mM sodium acetate were added for electron donors of *D. desulfuricans* and *G. sulfurreducens*, respectively. After 24-h incubation, cells were harvested by centrifugation at 10,000×g for 3 min. Whole mounts of the incubated cells of *D. desulfuricans* and *G. sulfurreducens* were observed by TEM (JEOL JEM-2010). The incubated cells *of D. desulfuricans* were embedded in LR White resin as described above, and 100-nm thick sections prepared with an ultramicrotome (Ultracut S, Reichert-Nissei, Tokyo, Japan) were also observed by TEM.

### Mineralogical and microbiological characterizations of powdered chimney samples

To clarify mineral compositions of the powdered subsamples, X-ray diffraction (XRD) pattern analysis was performed by Rigaku RINT 2000 powder diffractometer at 40 kV and 30 mA. The total cell numbers of the chimney interior and exterior were measured by a direct count method with SYBR Green I. The formaline-fixed powdered samples were suspended in 1:1 ethanol / phosphate buffered saline (PBS) solution. Some portions of the suspensions were sonicated for 30 sec. A 0.22-μm-pore-size, 25-mm-diameter polycarbonate filter (Millipore) was used to collect the sonicated suspension. To stain microbial cells, the filter was incubated in TAE buffer containing SYBR Green I for 5 min at room temperature. The stained filter was briefly rinsed with deionized water and observed under epifluorescence using the Olympus BX51 microscope with the Olympus DP71 CCD camera.

### DNA extraction and 16S rRNA gene amplicon analysis

Prokaryotic DNA was extracted from 0.1 g of frozen powdered chimney subsamples as described previously^45^. In brief, the powdered chimney subsample was incubated at 65°C for 30 min in 300 μL of alkaline solution consisting of 75 μL of 0.5 N NaOH and 75 μL of TE buffer (Nippon Gene Co.) including 10 mM Tris–HCl and 1 mM EDTA. After incubation, the aliquots were centrifuged at 5,000×g for 30 sec. Then, the supernatant was transferred into a new tube and neutralized with 150 μL of 1 M Tris–HCl (pH 6.5; Nippon Gene Co.). After neutralization, the DNA-bearing solution (pH 7.0–7.5) was concentrated using a cold ethanol precipitation, and the DNA pellet was dissolved in 50 μL of TE buffer. The purified DNA solution was stored at −4°C or −20°C for longer storage. For the negative control, DNA extraction from the subsample was performed in parallel with one extraction negative control to which no sample was added.

The 16S rRNA gene was amplified using LA Taq polymerase (TaKaRa-Bio, Inc.). For pyrosequencing, the 454 GS-junior sequencer (Roche Applied Science) was used. The primers Uni530F and Uni907R^46^ were extended with adaptor sequences (CCATCTCATCCCTGCGTGTCTCCGACTCAG for Uni530F and CCTATCCCCTGTGTGCCTTGGCAGTCTCAG for Uni907). The forward primer Uni530 was barcoded with 8-mer oligonucleotides^47^. Thermal cycling was performed with 35 cycles of denaturation at 95°C for 30 sec, annealing at 58°C for 45 sec, and extension at 72°C for 1 min. A PCR product with the expected size was excised from 1.5% agarose gels after electrophoresis on TAE (40 mm Tris acetate, 1 mm EDTA, pH 8.3), which was purified using MinElute Gel Extraction Kit (Qiagen, Inc.). A DNA concentration of the purified PCR product was measured by the Quant-iT dsDNA HS assay kit and the Qubit fluorometer (Invitrogen, Inc.). The concentration of total double-stranded DNA in each sample was adjusted to 5 ng/μL. Emulsion PCR was performed using the GS FLX Titanium emPCR Kit Lib-L (Roche Applied Science) to enrich DNA library beads for the 454 GS-junior sequencers. Amplified DNA fragments were sequenced according to the manufacturer’s instructions.

Raw reads were demultiplexed, trimmed and filtered based on their 8-bp sample-specific tag sequences, quality values (QV) and lengths using Mothur v. 1. 31^48^ to obtain unique reads more than 250 base pairs (bp), and an average quality score >27. Filtered sequences were aligned with Mothur to the Greengenes reference database^49^, and chimeric sequence reads were removed with Chimera Uchime in Mothur. Sequence reads were clustered into operational taxonomic units (OTU) sharing 97% identity within each OTU. Phylogenetic affiliations of the OTUs were assigned by the neighbor joining method in the ARB software package^50^, along with closely related sequences retrieved from GenBank (http://www.ncbi.nlm.nih.gov/genbank/) through BLASTn searches (somewhat similar sequences).

### Genome-resolved metagenomic analysis

From the subsample of the chimney interior, genomic DNA was extracted by a using an UltraClean Soil DNA Isolation Kit (MoBio Laboratories). As described previously^51^, shotgun library construction was performed for the extracted DNA using a KAPA Hyper Prep kit for Illumina (KAPA Biosystems), and sequencing of the library was performed on an Illumina MiSeq platform (MiSeq PE300). The trimming and filtering of the reads were performed using CLC Genomics Workbench (QIAGEN Aarhus A/S)^16^. After the high-quality reads were assembled using SPAdes^52^, the produced contigs were binned into a near-complete genome using Metabat^53^. The gene prediction and annotation of the contigs and the near-complete genome were annotated using Prokka^54^ and the Kyoto Encyclopedia of Genes and Genomes (KEGG) pathway tool ^55^ with the Blast-KOALA tool^56^. Initial analyses of near-complete genomes from Crystal Gyser^15^ were performed by ggKbase (https://ggkbase.berkeley.edu). For selected functional genes, closely related sequences and their source organisms were initially accessed using NCBI Protein BLAST Program^57^. Amino-acid sequences were aligned using Muscle^58^. To construct a neighbor-joining tree, the ARB software package was used^50^.

### Fluorescence *in-situ* hybridization analysis

Whole-cell hybridization was performed for thin sections of the intact chimney subsample embedded in LR-White resin. Hybridization was conducted at 46°C in a solution containing 20 mM Tris–HCl (pH 7.4), 0.9 M NaCl, 0.1% sodium dodecyl sulfate, 30% (v/v) formamide and 50 ng/μL of each probe labeled at the 5’ end with fluorescence dye. A Cy-5 labeled probe targeting the domain Archaea (Arch915: 5’-GTGCTCCCCCGCCAATTCCT-3’)^59^ and a Cy-5 labeled probe targeting Pacearchaeta (Pace915: 5’-GTGTCTCCCCGCCAATTCCT-3’) were used for hybridization of the positive control and the chimney thin sections, respectively. After hybridization, the specimens were washed at 48°C in a solution lacking the probes and formamide at the same stringency, adjusted by NaCl concentration. After staining with 4’,6-diamidino-2-phenylindole (DAPI) at 0.4 μg/ ml, the slides were examined using the Olympus BX51 microscope. For the positive control, cultured archaeal cells of *Methanocaldococcus* sp. Mc-365-70 were used. For the negative control, a bacteria-specific probe named EUB338 (5’-GCT GCC TCC CGT AGG AGT-3’)^60^ was used to check the absence of non-specific biding of the EUB338 probe on the cultured archaeal cells under the same hybridization conditions.

### DNA binding experiments to amorphous silica

A silica-based spin column was used to bind genomic DNA extracted from *Escherichia coli*(NBRC#20004). *E. coli* was grown at 30 in LB liquid medium and harvested by centrifugation at 8,000×g before reaching the stationary growth stage. The *E. coli* cells were subjected to DNA extraction as described above for metagenomic analysis. Some portions of the DNA extract diluted in seawater and TE buffer were loaded on silica-based spin columns (MoBio Laboratories). After loading of the diluted DNA extracts, the silica-based spin columns were centrifuged at 10,000×g to recover the loaded DNA extracts. DNA concentrations in the DNA extracts before and after loading to the silica-based spin columns were measured with by the Quant-iT dsDNA HS assay kit and the Qubit fluorometers. For the seawater-based experiment, the DNA-loaded spin column was eluted with TE buffer. The recovered DNA was quantified as described above. Fourier transformed infrared-ray (FT-IR) spectrometry (Shimadzu IRSprit) was used to clarify the crystallinity of silica. Powder materials were analyzed by a mode of attenuated total reflection (ATR) through KBr beam-splitter with temperature-controlled DLATGS detector. Representative FT-IR spectra were collected by averaging 45 individual spectra. Amorphous silica (Aldrich) was used for a reference.

## Supporting information

https://sendfile.s.u-tokyo.ac.jp/public/5jYoQACJw8DA2R4BlvV40JwVXSB3YAuPrcS0WeKgWPjp

https://sendfile.s.u-tokyo.ac.jp/public/gjZMQATJCYDALW8BHXx4z5QQTnFQ9nrK_sSe9MWR0Rvb

## Data and Code availability

All data needed to evaluate the conclusions in the paper are present in the paper and/or the Supplementary Materials. The 16S rRNA gene sequences in this study were all deposited in the DDBJ nucleotide sequence database with accession numbers LC554901-LC555740. The metagenome-assembled genome sequences were deposited with accession numbers BHXO01000001–BHXO01000035 under the BioProject accession number, PRJDB6687. The Pacearchaeota genome constructed in this study was named as IPdc11 in the database.

## Acknowledgements

We thank captain and crews of the R/V Natsushima and the ROV Hyper-Dolphin operating groups for their technical support in sample collection. We also thank to the scientists that joined the NT12-24 cruise and to the members of TAIGA project for providing opportunity of this study. We are grateful to Koji Ichimura and Miho Hirai for their technical assistance. We are also grateful to Kohei Kitamura for the arrangement of iMScope, and Satoko Nishikawa, Yuka Soma and Morihiko Onose for operating Raman spectroscopy. We thank Edanz Group (https://en-author-services.edanz.com/ac) for editing a draft of this manuscript

