## Supplementary material for "Ultra-small cells and DPANN genome unveiled inside an extinct vent chimney": https://sendfile.s.u-tokyo.ac.jp/public/5jYoQACJw8DA2R4BlvV40JwVXSB3YAuPrcS0WeKgWPjp

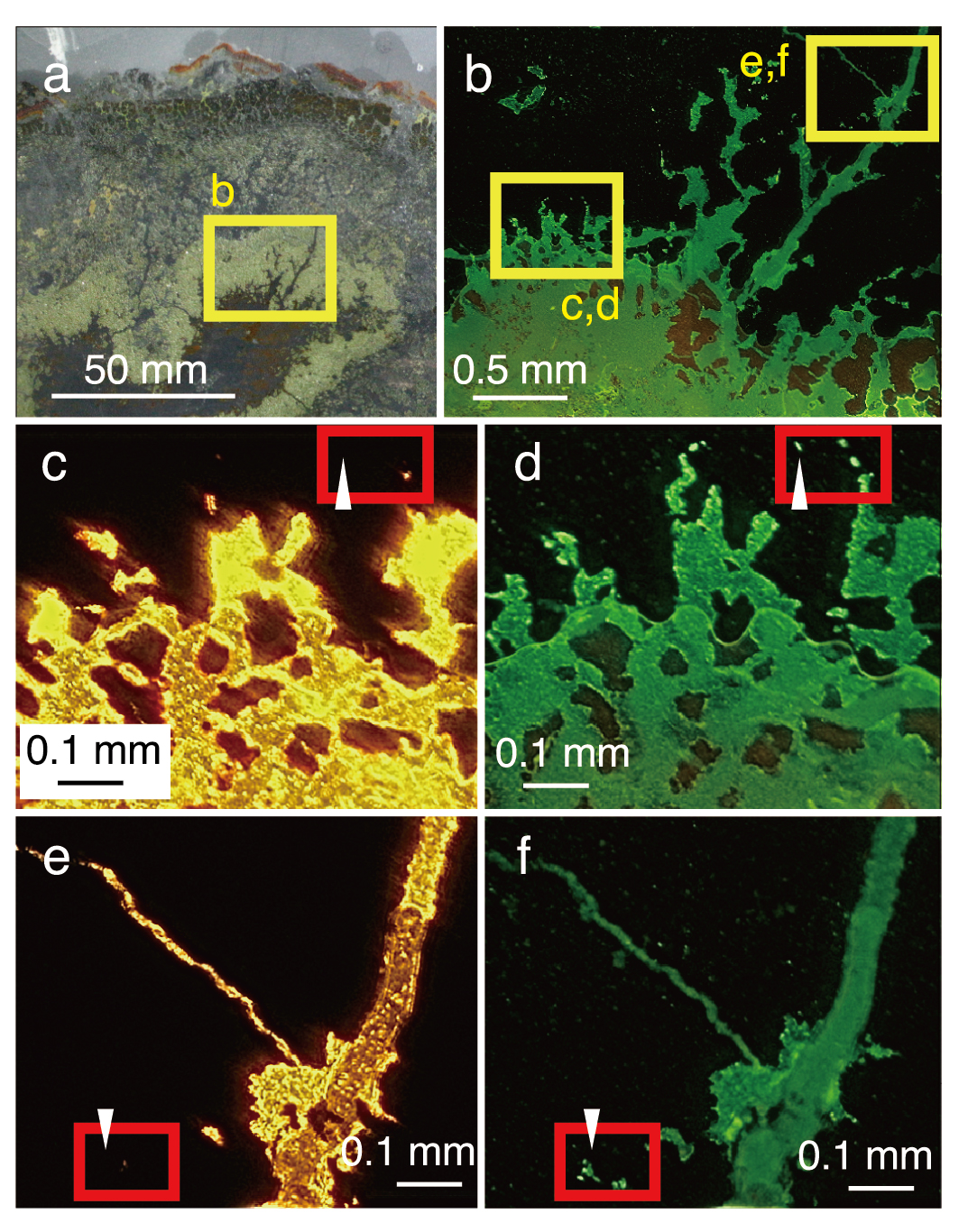

**Supplementary Figure 1| Light and fluorescence microscopy images of a SYBR Green I-stained thin section from the extinct chimney.** **a,** Optical microscope image. **b,** Low-magnification fluorescence image. **c,** Middle-magnification transmitted-light image. **d,** Middle-magnification fluorescence image. **e,** Middle-magnification transmitted-light image. **f,** Middle-magnification fluorescence image. Yellow rectangles highlight areas observed at higher magnifications in separate images. Red rectangles and arrows highlight SYBR-Green stained grain boundaries without light transmission.

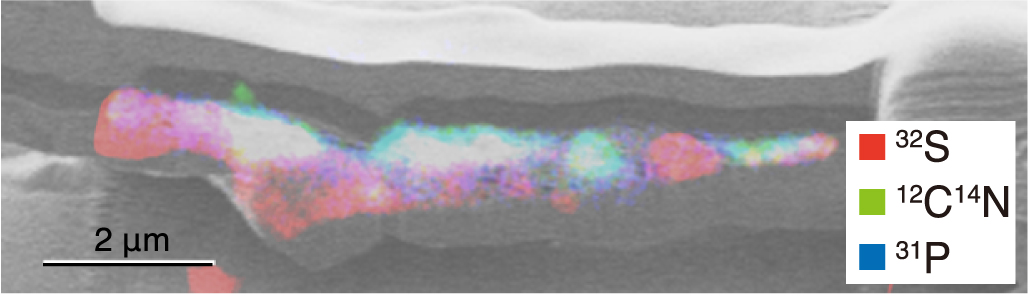

**Supplementary Figure 2| Elemental distribution in a silicate-bearing layer with sub-micron voids.** Overlay image of a Ga ion image in black and white and the Nanoscale secondary ion mass spectrometry (NanoSIMS) images of ^32^S in red, ^12^C^14^N in green, and ^31^P in blue.

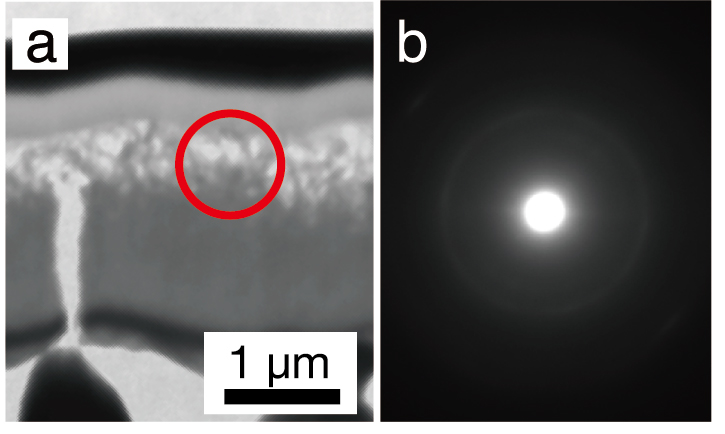

**Supplementary Figure 3| Mineralogical characterizations of a fibrous silica-bearing layer in a 300-nm thick FIB section.** **a,** Transmission electron microscope (TEM) image. **b,** A selected area electron diffraction (SAED) pattern. Supplementary Figure 3b was taken from a red circle in Supplementary Figure 3a.

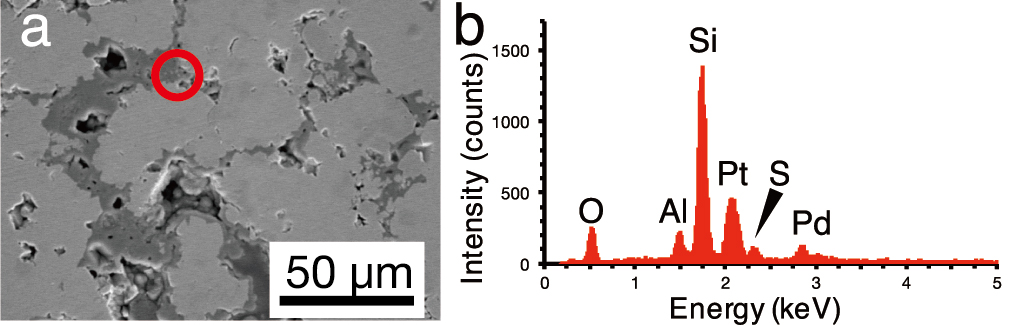

**Supplementary Figure 4| Mineralogical characterizations of a region fabricated for a 150-nm thick FIB section. a,** Scanning electron microscope (SEM) image. **b,** Energy-dispersive X-ray spectroscopy (EDS) spectrum of a red circle in Supplementary Figure 4.

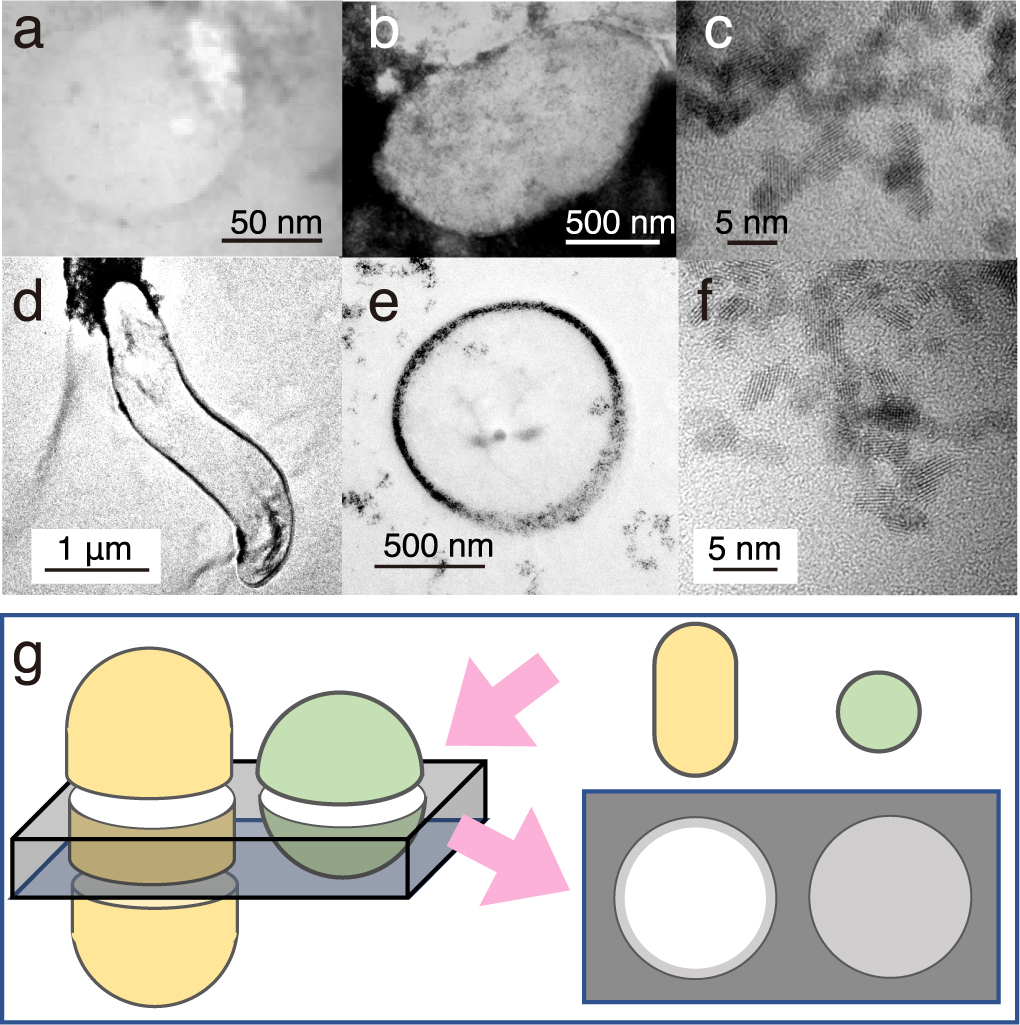

**Supplementary Figure 5| TEM image contrast from microbial cells associated with extracellular nanoparticles. a,** Typical small sphere observed in Figure 2e. **b-c,** *Geobacter sulfureducens* extracelluarly precipitated with uraninite nanoparticles. **d-f,** *Desulfovibrio desulfruicans* with periplasmic uraninite nanoparticles**. g,** Schematic explanation of the appearance of microbial cells extracellularly coated with nanoparticles by TEM observations. This illustration shows how a 150-nm thick FIB section was fabricated for rod and coccoid cells (left) and an expected TEM image from the FIB section (right). Gray color indicates the presence of extracellular nanoparticles in the right illustration.

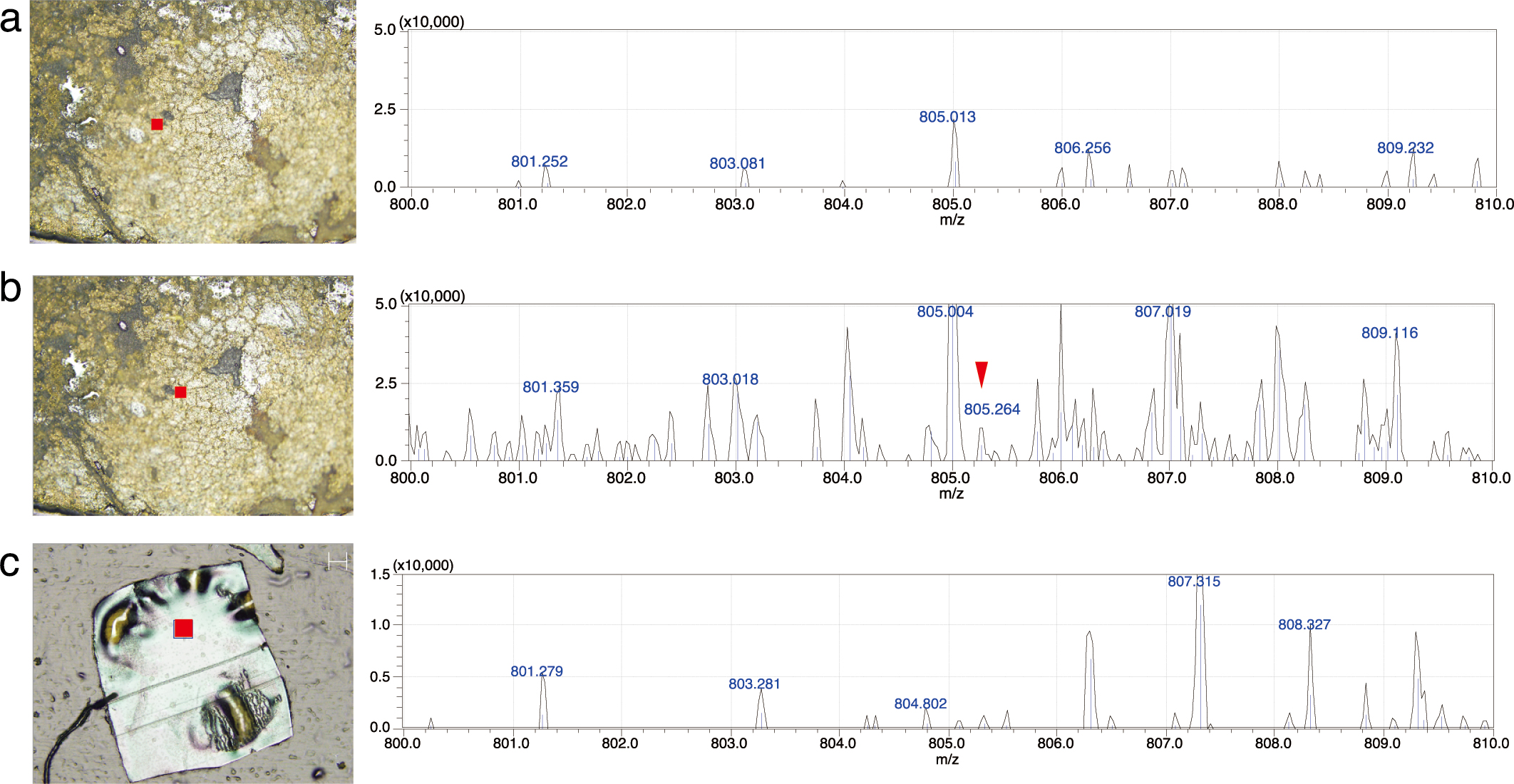

**Supplementary Figure 6| Imaging mass spectrometry analysis of grain boundaries of the inner chalcopyrite wall and the resin used for embedment.** Spot analysis was conducted at each filled square in the left images. **a-b,** Chalcopyrite grain boundary. **c,** Resin in a 5-μm-thick section.

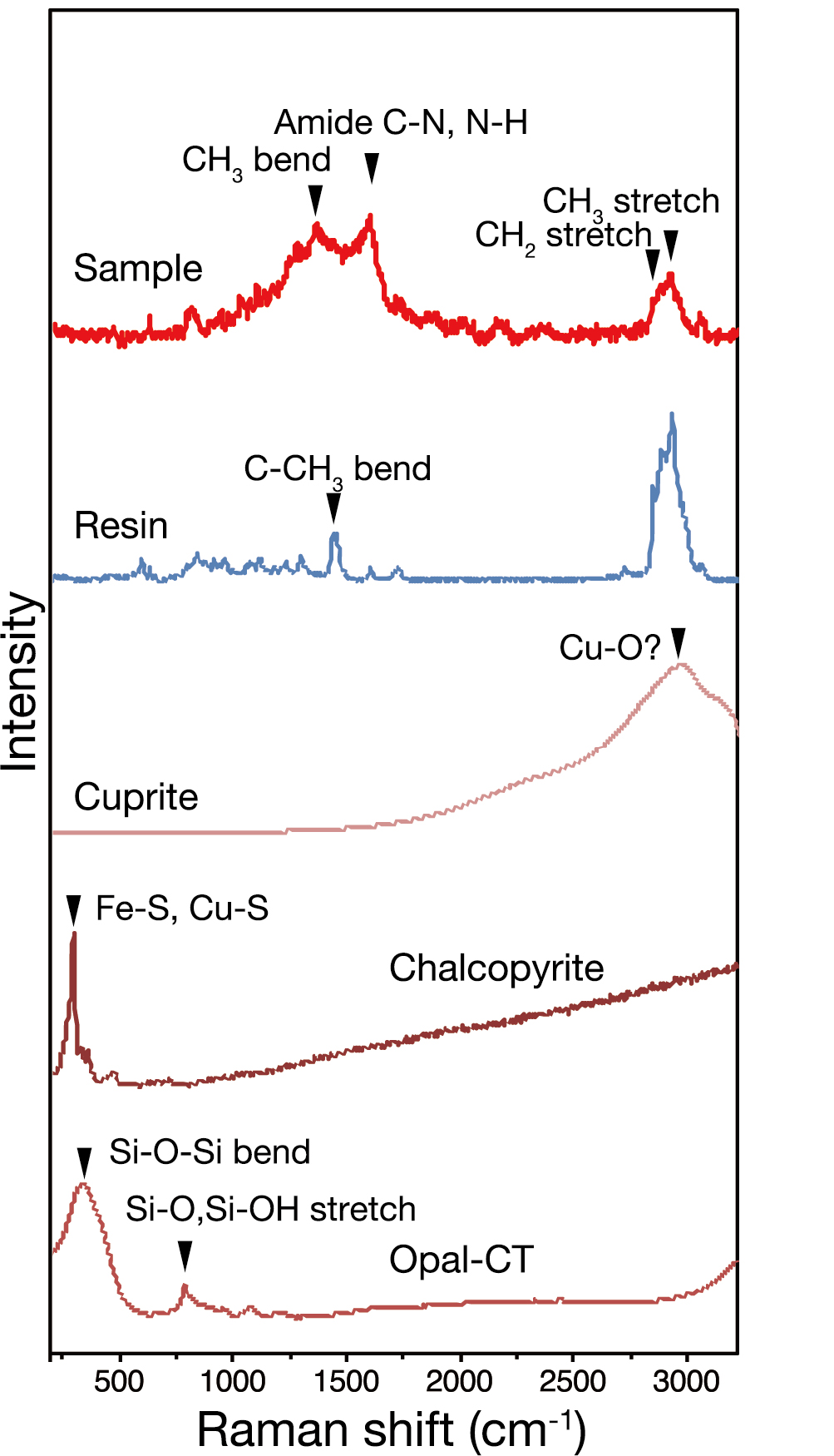

**Supplementary Figure 7|** **Comparison of Raman spectra from a grain boundary in the inner chalcopyrite wall and references.** A spectrum marked with “sample” is also shown in Fig. 4b. A spectrum marked with “resin” was obtained from LR White resin. Cuprite, chalcopyrite and opal-CT were shown as spectra obtained from RRUFF (<http://rruff.info)>^1^. Peak positons in opal-CT are almost the same as those in amorphous silica^2^.

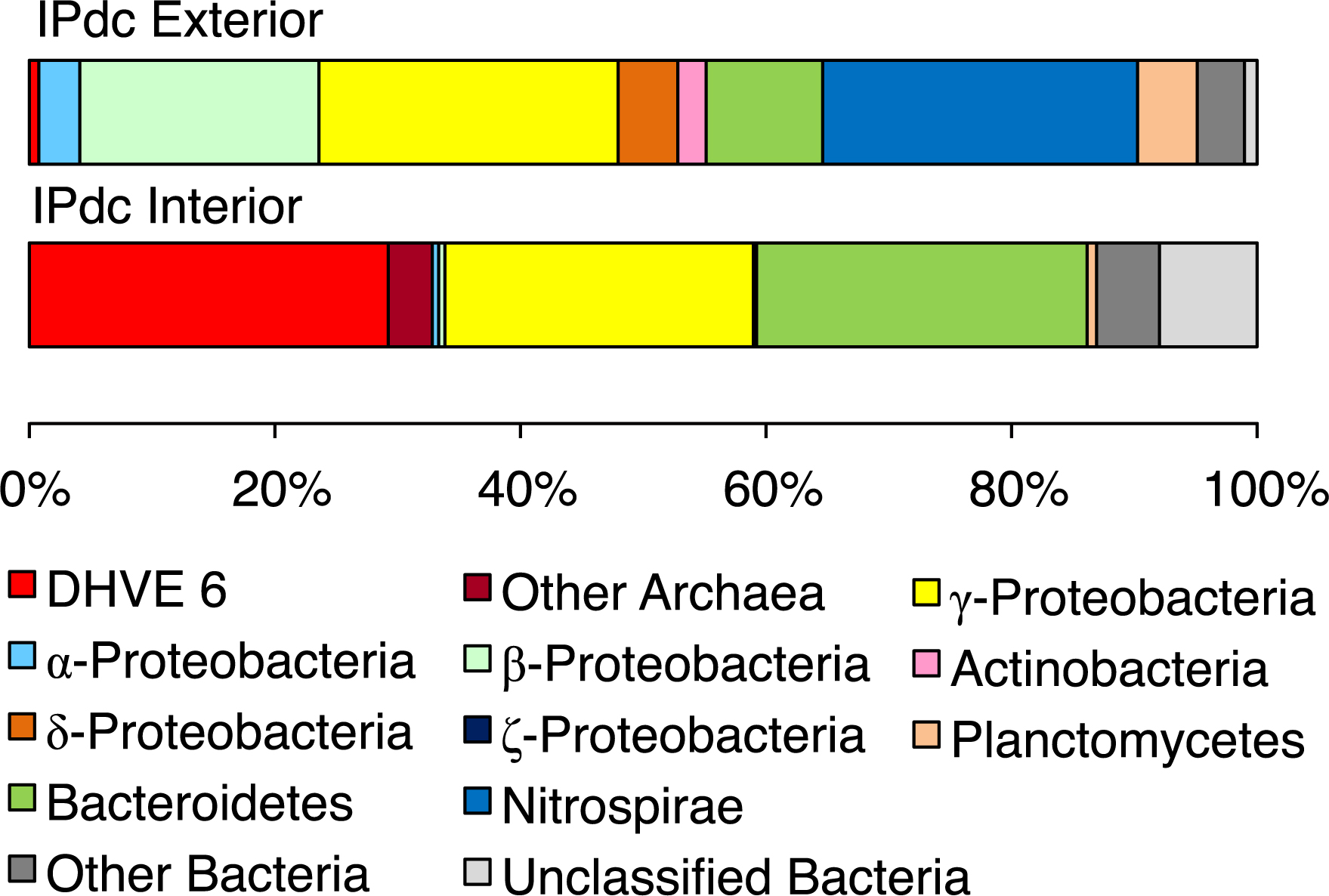

**Supplementary Figure 8| Microbial community structures based on prokaryotic 16S rRNA gene sequences from the chimney interior and exterior.** The relative abundances of sequences classified by phylum or class are shown.

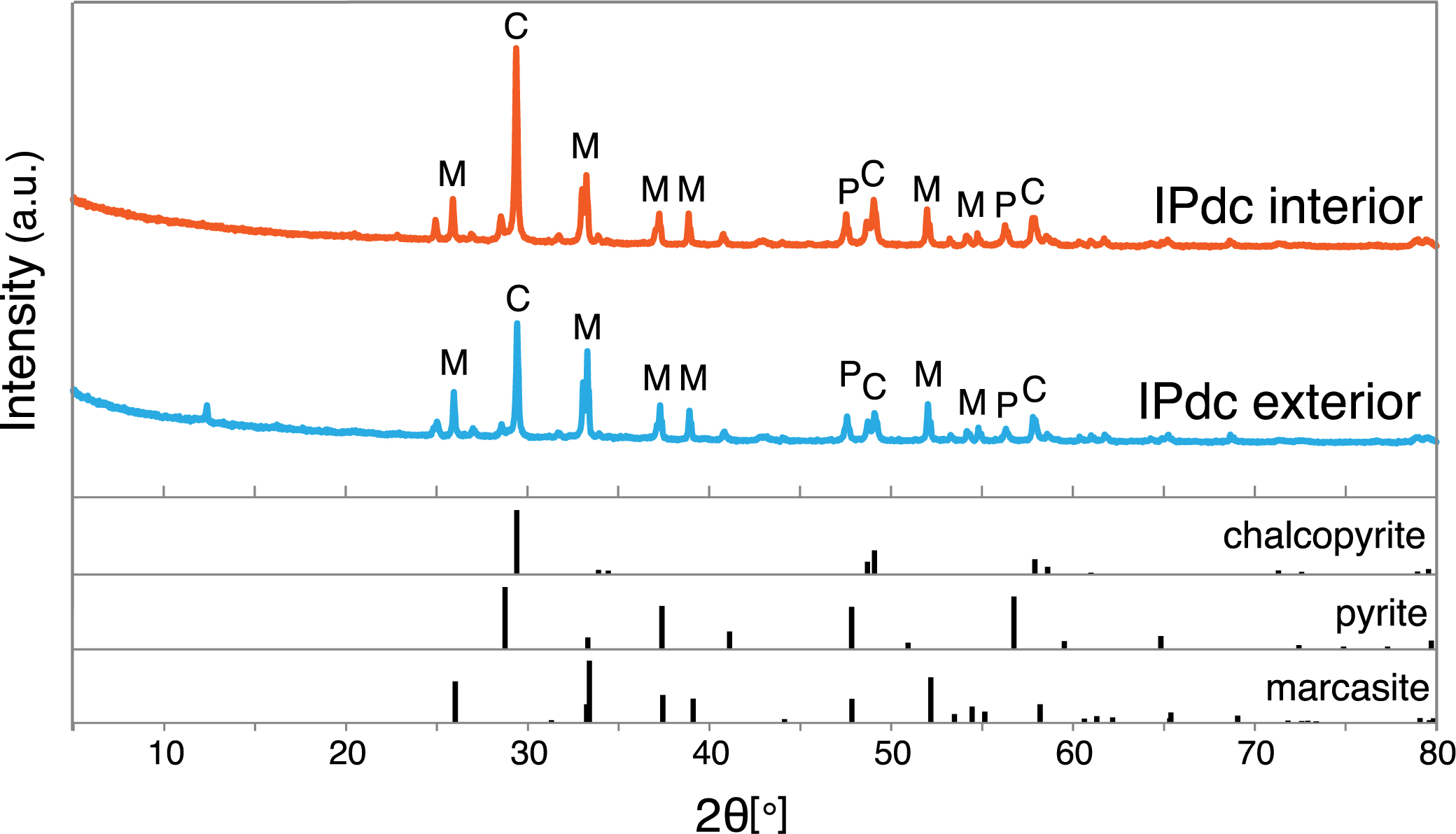

**Supplementary Figure 9| Mineral assemblages in the chimney interior and exterior.** Powder XRD patterns of the extinct chimney and those from minerals from American Mineralogist Crystal Structure Database)^3^. Peaks from chalcopyrite, pyrite, and marcasite are labeled with C, P, and M, respectively.

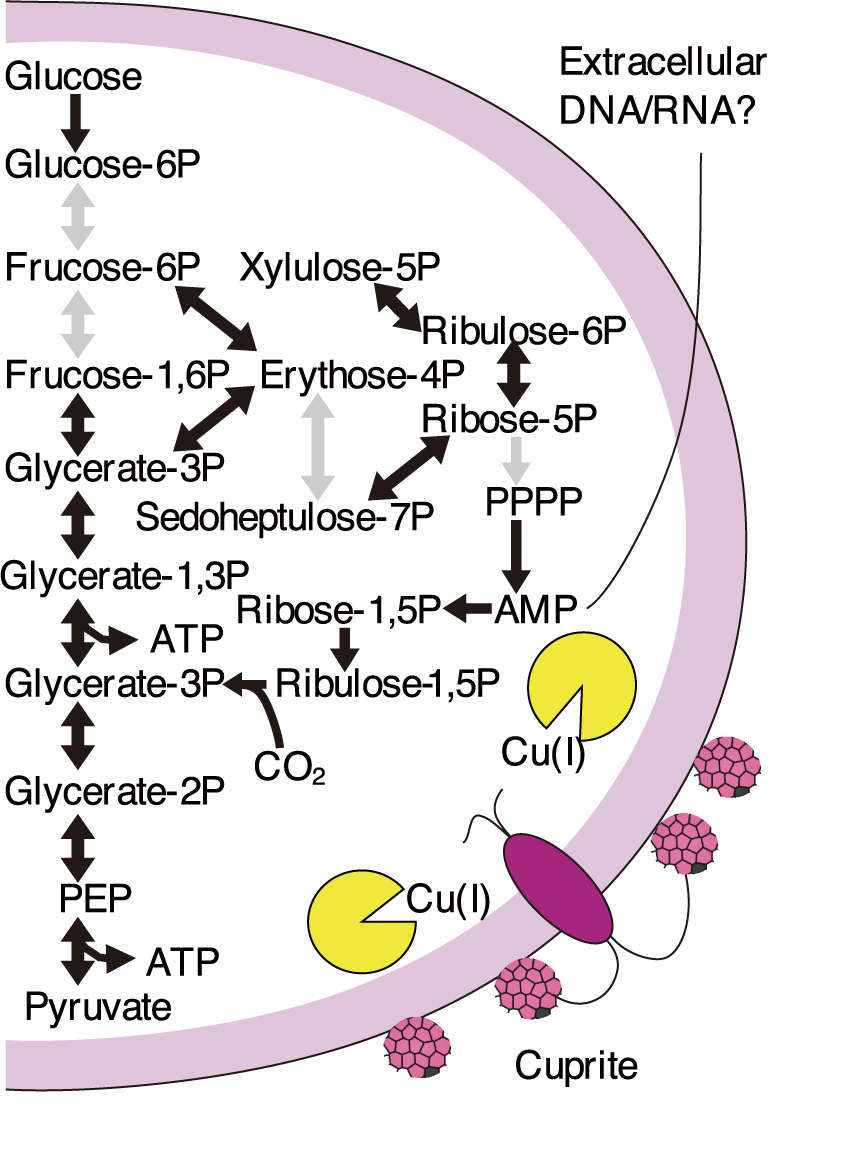

**Supplementary Figure 10**| **Schematic illustrations of inferred metabolic pathways** Pathway map showing the carbon metabolism with black and gray arrows indicating the presence and lack of genes and the copper exporting system (lower right). Black lines indicate cellular uptake and excretion of metabolites. The lack of genes does not indicate their absolute absence in the original genome, because of the incompleteness of genome reconstruction.

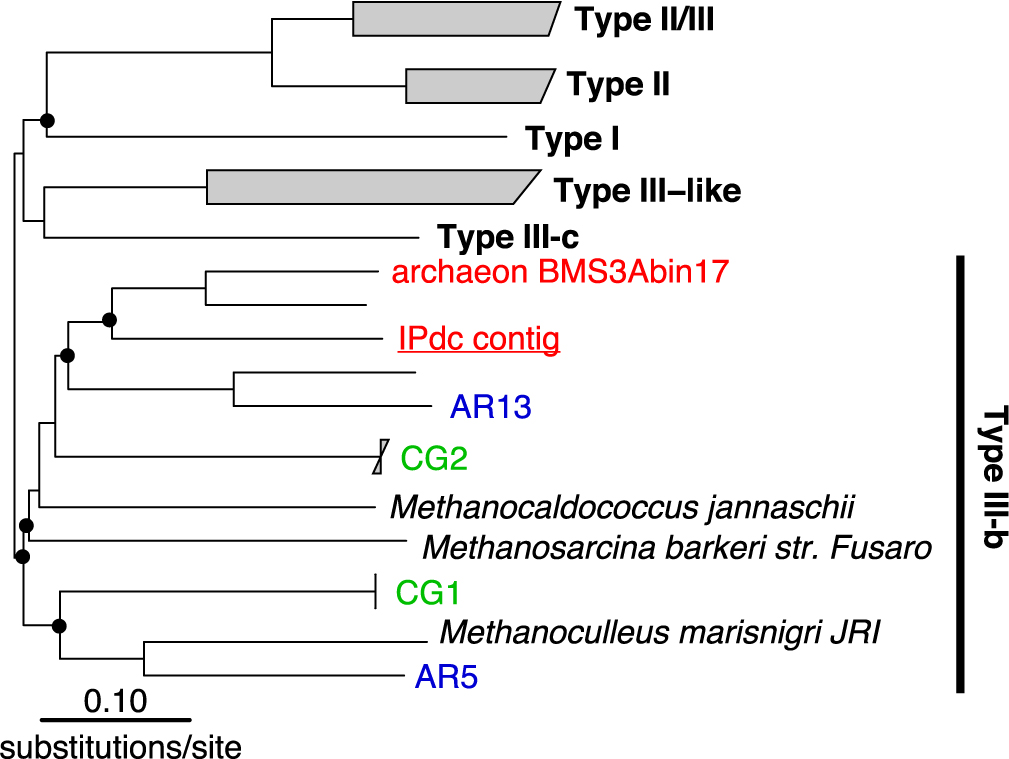

**Supplementary Figure 11**| **Phylogenetic classification of RuBisCO.** Neighbor-joining phylogenetic tree of amino acid sequences from RuBisCO TypeIII-b. AR and CG Pacearchaeota genomes are shown in blue and green, whereas IPdc and BMS3Abin17 Pacearcheota genomes are shown in red. Branches with bootstrap values >50% are indicated with filled circles.

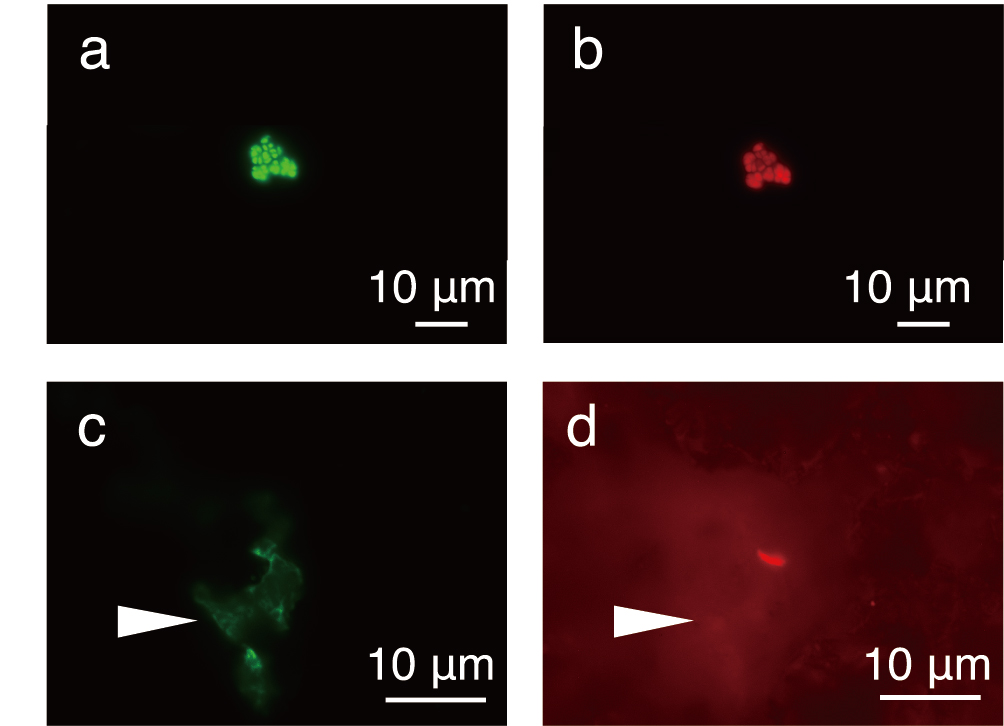

**Supplementary Figure 12| Application of FISH to resin-embedded microbial cells in thin sections.** **a-b**, Cultured cells of *Methanocaldococcus* sp. Mc-365-70 stained with SYBR-Green I (left) and a Cy5-labelled Archaea-universal probe called Arch915 (right). **c-d**, Grain boundary of the inner chalcopyrite wall stained with SYBR-Green I (left) and a Cy5-labelled Pacearchaeota-targeted probe called Pace915 (right). A white arrow indicates a grain boundary region, in which greenish spots stained by SYBR-Green I (left) are not associated with Cy5-derived signals (right).

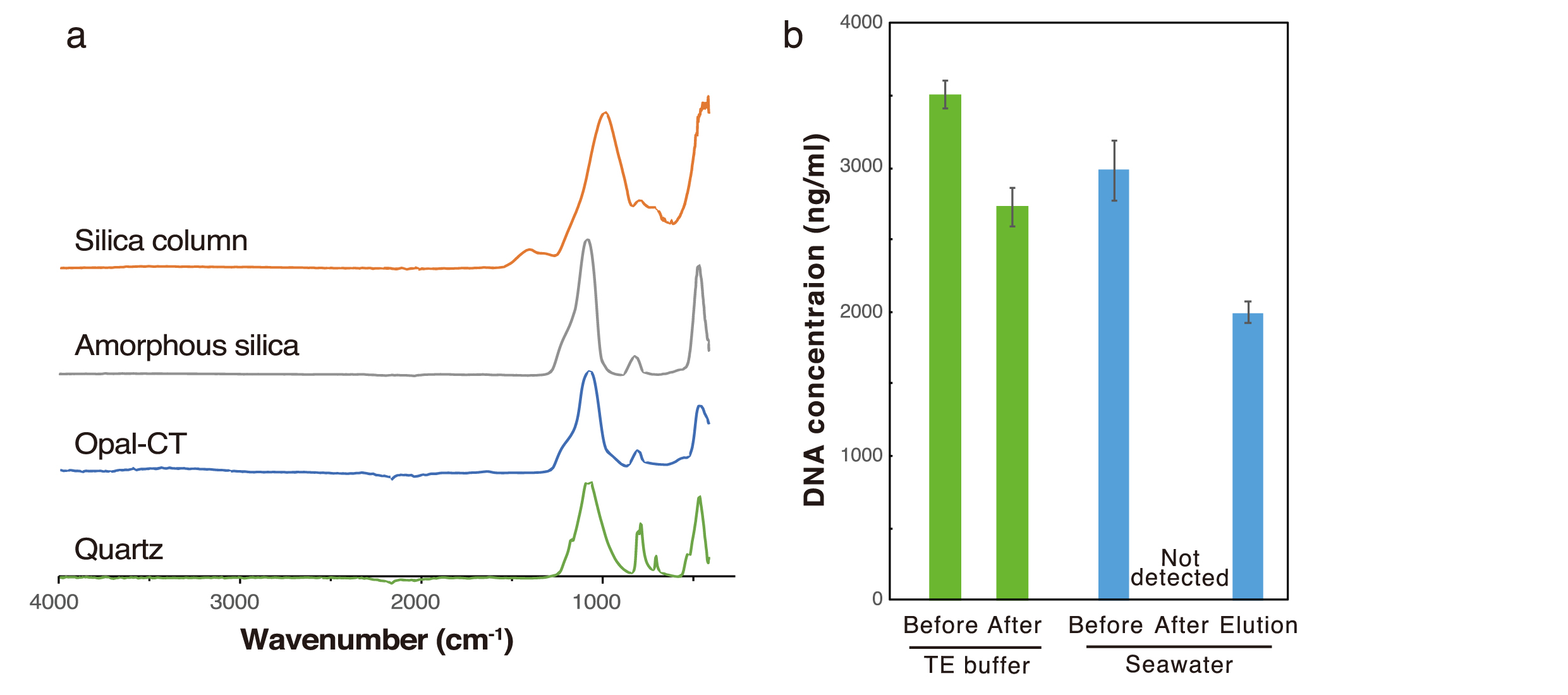

**Supplementary Figure 13|** **Ability of amorphous silica to bind DNA from seawater.** Silica-based spin columns were used to examine DNA binding in TE buffer (left green) and seawater (right blue). **a**, Silica crystallinity evaluated by Fourier transformed infrared-ray (FT-IR) spectrometry. FT-IR spectra of silica used in a spin-column and reference spectra from amorphous silica and RRUFF database^1^. **b,** DNA concentrations measured before and after loading into the spin columns. For the seawater-based experiment, TE buffer was used for elution of column-trapped DNA after loading from seawater.

1	Lafuente, B., Downs, R., Yang, H. & Stone, N. The power of databases: the RRUFF project. In “Highlights in mineralogical crystallography”, Armbruster, T. & Danisi, RM, eds. W. *De Gruyter, Berlin, Germany* **1**, 30 (2015).

2	Sodo, A. *et al.* Raman, FT‐IR and XRD investigation of natural opals. *J. Raman Spectrosc.* **47**, 1444-1451 (2016).

3	Downs, R. T. & Hall-Wallace, M. The American Mineralogist crystal structure database. *Am. Mineral.* **88**, 247-250 (2003).
